# Reconstructing spatial transport distributions in the nuclear pore complex from 2D images—how reliable is it?

**DOI:** 10.1101/145110

**Authors:** Li-Chun Tu, Maximiliaan Huisman, Yu-Chieh Chung, Carlas Smith, David Grunwald

## Abstract

Imaging single molecules in living cells and reconstituted cell systems has resulted in a new understanding of the dynamics of nuclear pore complex functions over the last decade. It does, however, fall short on providing insights into the functional relationships between the pore and nucleocytoplasmic cargo in three-dimensional space. This limited ability is the result of insufficient resolution of optical microscopes along the optical axis and limited fluorescent signal due to the short timescales involved in nuclear transport (fractions of a second). To bypass current technological limitations, it was suggested that highly time-resolved 2D single molecule data could be interpreted as projected cargo locations and could subsequently be transformed into a spatial cargo distribution by assuming cylindrical symmetry ^1^. Such cargo distributions would provide valuable insights into the NPC-mediated transport in cells. This method, termed 3D-SPEED, has attracted large interest inside and beyond the nuclear pore field, but has also been sharply critiqued for a lack of critical evaluation. Here we present such an evaluation, testing the robustness, reconstruction quality and model-dependency.

---

Nuclear pore complexes (NPCs) span the nuclear envelope and mediate bidirectional transport between nucleus and cytoplasm. Macromolecules (>40 kDa) require transport receptors to transit the NPC efficiently, whereas smaller molecules diffuse through the NPC passively^1^. As such, NPCs play an essential role in maintaining cellular homeostasis and preventing viral replication. While the scaffold-structure and composition of the NPC have been resolved in great detail^2^, the mechanism by which the semi-permeable barrier at the center of the NPC regulates selective transport is unknown. Aiming to elucidate this mechanism, Ma, J., Goryaynov, A., and Yang, W. (NSMB 2016;23, 239-247) set out to investigate spatial cargo distribution within the nuclear pore complex in their manuscript ‘Super-resolution 3D tomography of interactions and competition in the nuclear pore complex’ using so-called *SPEED* (single-point edge-excitation subdiffraction) microscopy^3–7^.

*SPEED* microscopy features innovations in optical microscopy as well as data processing, and has been the topic of several review articles ^8–12^. The potential of *SPEED* to obtain pseudo-tomographic data prompted us to systematically analyze the technical aspects of the data processing method. Through a series of simulations, we examined the data requirements (precision, size of dataset, symmetry constraints, etc.) and the limits to which reconstructed transport densities can be interpreted with confidence (see Supplement).

The *SPEED* process uses two-dimensional (2D) single-molecule localizations obtained by high-speed fluorescence microscopy to reconstruct a three-dimensional (3D) density distribution. This back-projection transformation assumes that the transported particles are distributed in a cylindrically symmetrical manner, requiring just a single ‘perspective’ to reconstruct the spatial distribution. The symmetry constraint has two important implications: (1) if the underlying transport distribution is not fully cylindrically symmetric, then the reconstructed 3D density does not correspond to the actual 3D density; and (2) if the 2D localization data used to obtain the 3D density does not reflect the underlying cylindrical symmetry (i.e. due to under-sampling or imprecision), then the accuracy of the reconstructed 3D density will diminish. As such, 3D SPEED does not meet the definition of tomography and is not guaranteed to provide accurate or even unique reconstructions.

We simulated idealized localization datasets with known underlying distributions, performed the *SPEED* transformation, and then compared reconstructed distributions to the ground truth input distributions **(Fig. 1a-f)**. The resulting comparisons **(Fig. 1g-i)** revealed that the quality of reconstruction depends strongly on the size of the dataset and the overall precision of the transport particle localization: average success rates upwards of 75% typically require at least a few hundred localizations and a localization precision below 5nm for the distributions we simulated. Registration precision – chromatically and between separate data sets – the precision of detection of NPC rotation, and the degree of symmetry of the transport distribution also critically affected the quality of the reconstruction (see Supplement). Our results indicate that, under the reported experimental conditions (^~^100 localizations and 10-13nm localization precision) ^4–8,13,14^, the SPEED technique cannot reliably distinguish between uniform, central, peripheral, and bimodal transport in the central channel of the NPC, as claimed^7^. Increasing the size of a dataset and improving localization precision would improve the reliability of *SPEED,* but to do so would require precision enhancements that are currently out of reach in highly time-resolved live-cell optical microscopy.

**Figure 1.**
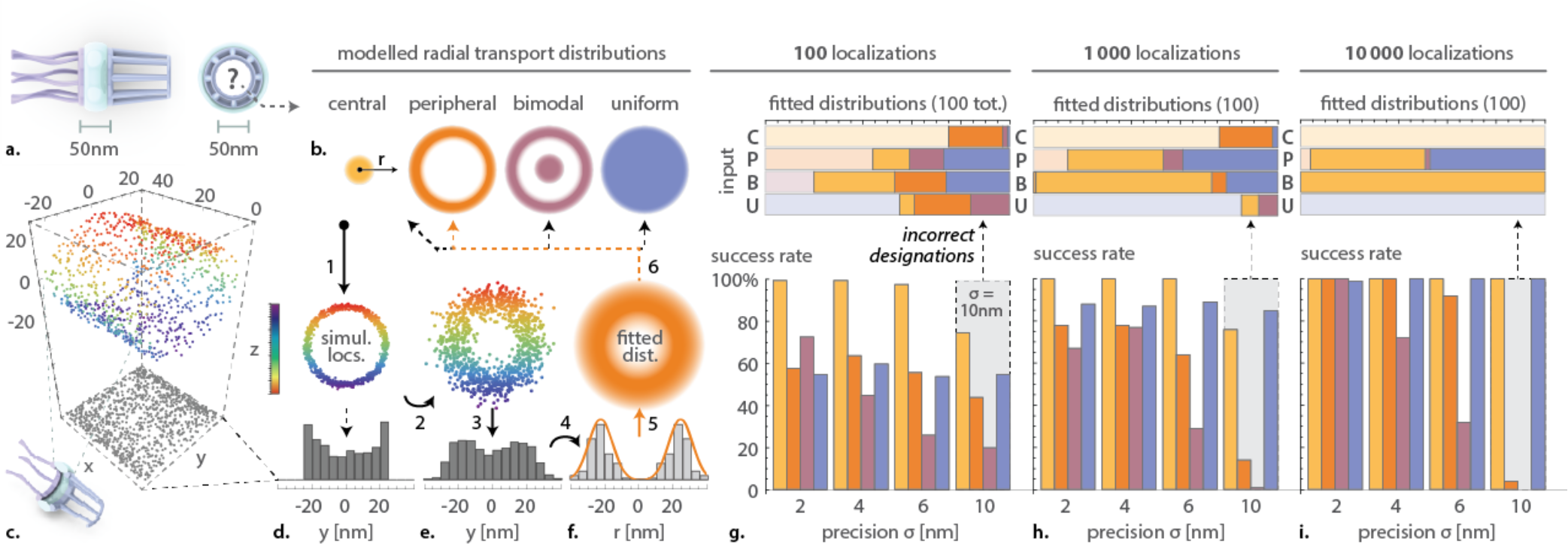
The performance of *SPEED* depends on the dataset size (number of localizations) and overall precision. ***SPEED*** was used to reconstruct radial cargo distribution in the central channel of the NPC using idealized, simulated data. (**a**) Dimensions of the NPC central channel. (**b**) The central channel accommodates transport of molecules through its center or periphery. (**c-f**) To quantify the ability of ***SPEED*** to distinguish between central, peripheral, bimodal, or uniform distribution, a simulated dataset of transport localizations was generated for each distribution pattern (step 1; shown in color in **c**). Simulated measurement imprecision was added (step 2), and then the localizations were projected to 2D (step 3; grey dots in **c**), to mimic data acquired by microscopy. The projected density profiles were extracted in **d** and **e** (note the effect of the simulated measurement imprecision in **e** compared to **d**). The back-projection transformation was subsequently applied (step 4) to the projected density profile in **e** to reconstruct a radial density profile in **f**, which was then fitted to a radial distribution (step 5) and compared to the four model distributions (step 6). The most significant fit was compared to the input distribution of the simulated dataset, revealing either successful identification or failure. (**g-i, bottom charts**) Success rates (% correctly identified) of 100 different simulations for each combination of distribution type (central, peripheral, bimodal and uniform), dataset size (10^2^, 10^3^, and 10^4^ localizations) and precision (σ = 2, 4, 6, 10 nm). (**g-i, top charts**) Identities of the incorrectly designated reconstructions (opaque colors) assigned by the ***SPEED*** data processing algorithm for each dataset size at precision σ = 10 nm reveal that reconstructions mainly fail due to under-sampling in small datasets and due to blurring (imprecision) in large datasets.

## Acknowledgements

This work was funded by grants 1R01GM123541-01 and 1U01EB021238 to D.G. We would like to thank Dr. S. Musser for critical feedback on the whole project and Mathias Hammer for critical reading of theory development.

## Author contributions

All authors contributed to the study design and data interpretation; LCT, MH and YCC collected and analyzed data, MH lead and implemented simulations with support from CS and YCC; YCC derived theory with support from LCT and CS. DG initiated and supervised project. All authors wrote, revised and approved the manuscript.

## Supplementary Information

### Introduction

The 3D-SPEED method as described by Yang et al. proposes a number of conceptual and experimental innovations, ranging from the image acquisition to data-processing and subsequent interpretation^1^. The technique led to a number of high impact primary publications ^1–5^ as well as numerous reviews, e.g. ^6–9^. Although a number of challenges have been pointed out pertaining to the acquisition side of the proposed microscopy technique (see discussion), our analysis focuses on the reconstruction of radial transport distributions for particles traversing the NPC using projected cargo positions obtained from idealized 3D-SPEED microscopy data. If we embrace the assumption that the spatial distributions of NPC cargo are cylindrically symmetrical around the central axis of the pore, these distributions should appear the same when looking from the side (z-direction in **Figure S1a)** - regardless of the perspective rotation relative to the NPC axis. Consequently, this assumption would allow an inverse projection transformation to translate a projected cargo distribution ((*x,y*)-plane in **Figure S1a)** to a more useful radial cargo distribution (in cylindrical coordinates **–** see **Figure S1ef)**. While the proposed reconstruction process is mathematically straight-forward (see subsequent chapter for an analysis), practical use of this method demands a number of considerations.

**Figure S1.**
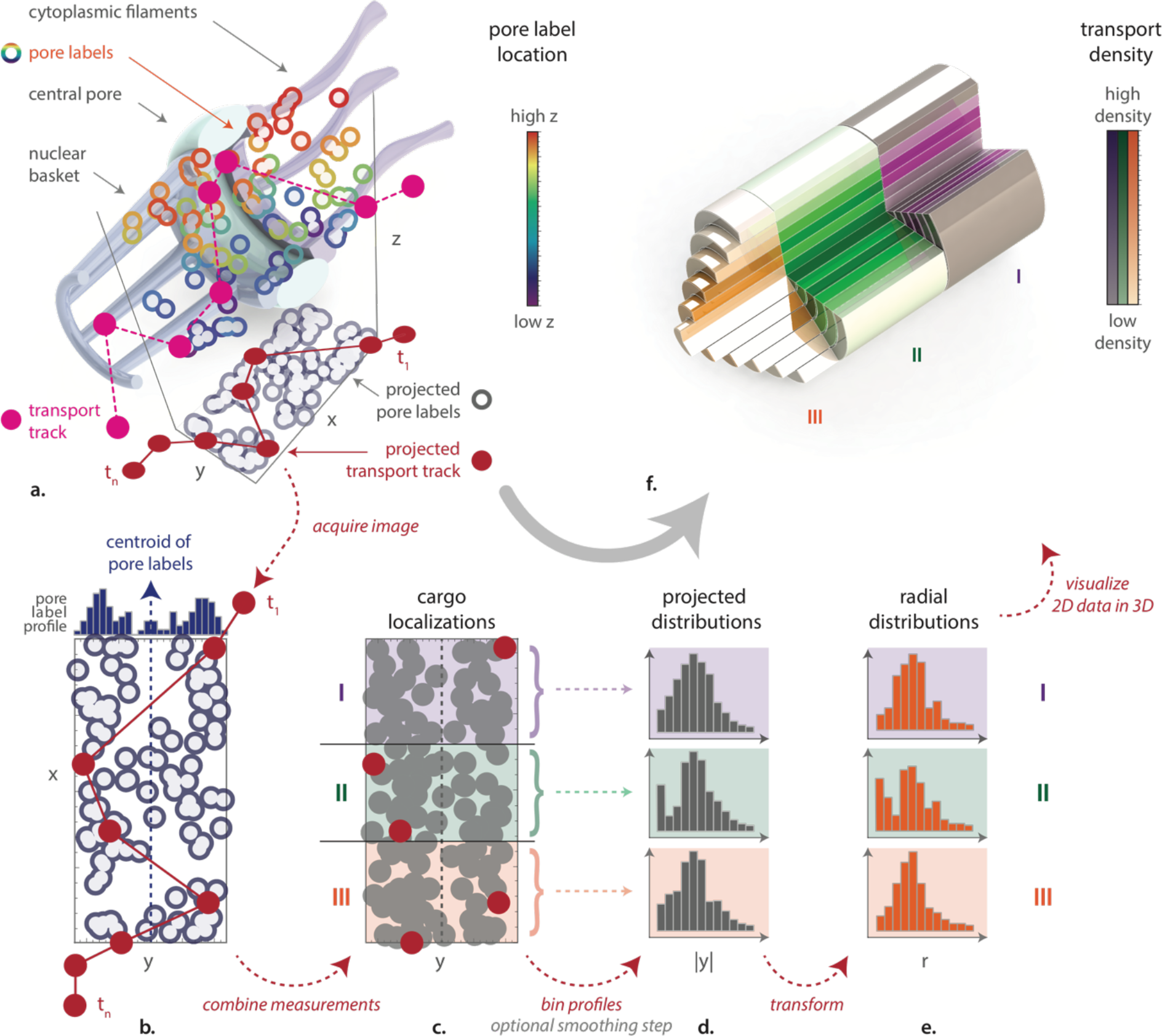
Reconstruction of radial NPC-cargo distribution from projected single molecule localizations. In two separate color channels, a projection of the fluorescently labelled nucleoporins (open circles) and cargo particles (closed circles, pink indicates 3D positions; red indicates projected 2D positions recorded on camera) are imaged using time-lapse fluorescence microscopy (**a**). Transport tracks from multiple measurements are combined through registration using the centroid of the nuclear pore labels (**b**; blue dashed line) to increase the number of cargo localization in the dataset (**c**). The dataset can be split into multiple segments along the transport axis to study how cargo densities are region-dependent (three segments are shown here in purple, green and orange). For each segment, a projected cargo distribution is obtained by binning along the x-axis (**d**).If axial symmetry is assumed, the profiles in both directions along the y-axis can be combined to obtain a profile |y|. By performing an inverse-projection transformation that assumes cylindrical symmetry, a radial distribution of the cargo particles can be calculated for each segment (**e**). A three-dimensional visualization of such a cargo distribution is shown in (**f**), where the color intensity depicts the reconstructed average cargo density in each segment of the central channel.

### Detailed results and discussion

The proposed reconstruction method has certain implications that need to be considered when interpreting the results. Firstly, a finite set of points cannot exhibit complete cylindrical symmetry, which necessitates the use of particle densities rather than spatial coordinates for reconstruction (see methods). This begs the question of how many cargo localizations are required to produce projected cargo densities that accurately reflect the underlying distribution. We investigate this question using simulated datasets of varying sizes **(Figure S2)**. Since these simulated datasets assume an idealized measurement system (e.g. zero localization imprecision, infinitely accurate registration, zero drift) it allows the requirements on the part of the dataset size to be evaluated independently of other experimental factors. One of the most striking observations was the occurrence of negative probabilities in the reconstructed distributions **(Figure S2c)**.

**Figure S2.**
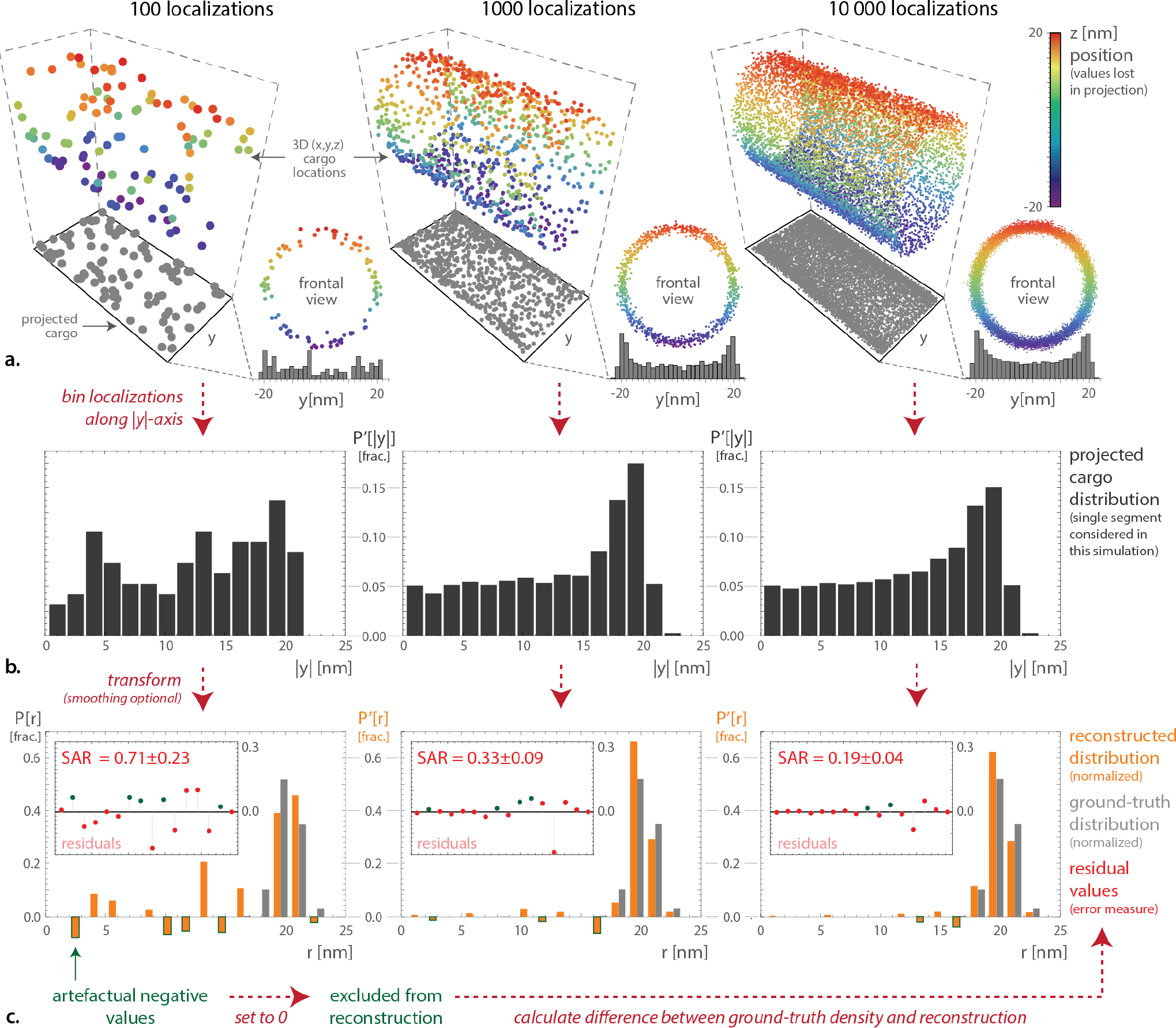
Accurate reconstruction of the radial cargo distribution requires a sufficient number of cargo localizations. Three 3D cargo localization datasets with 100 (left), 1000 (center) and 10000 (right) cargo particles were simulated (**a**; colored dots) with the same underlying radial distribution. Orthogonal projections (**a**; grey dots) were obtained by removing the z-coordinate, after which projected localization distributions were binned along the y-axis and mirror-summed assuming the central axis of the NPC was known (r=0; i.e. no centroid extraction was performed) (**b**). After applying the inverse-projection transformation, reconstructed radial cargo distributions (**c**; orange bars) are compared to the ground-truth distribution (**c**; grey bars) to yield residual plots (**c**; insets). For small datasets (left), artefactual negative probabilities are frequently found in the reconstructed distribution (green outlines); these values are set to zero (affecting some of the residual values as indicated by the green dots), after which the distributions are normalized for comparison. The SAR values in the insets (mean±SD) represent the sum of absolute valued residuals for 15 independent simulations and serve as an error-measure to compare reconstruction quality.

Since these negative probabilities largely disappear as the size of the dataset (number of particle localizations) increases, we conclude that these are artifacts arising from the fact that a limited number of cargo localizations cannot represent a cylindrically symmetrical distribution. After excluding negative probabilities by setting them to zero and normalizing the resulting distribution, we calculate the sum of the absolute-valued residuals (SAR), which serves as an error-measure for the mismatch between the ground-truth and the reconstructed transport distribution (see methods section). By repeating this process for independently simulated cargo localization datasets, the average reconstruction quality, as well as its variability, can be quantified using the average SAR value and its standard deviation, respectively. Simulating a range of dataset sizes **(Figure S3a)** quantitatively reveals a trend shown in **Figure S2c**: reconstruction quality critically depends on the dataset size. While smoothing the data before transformation can improve the reconstruction **(Figure S3b)**, its effect is limited and distribution-dependent (data not shown). The shape of the distribution also affects the reconstruction quality: as we expected considering the symmetry condition, narrow distributions require fewer data points to be reconstructed accurately **(Figure S3c)**.

**Figure S3.**
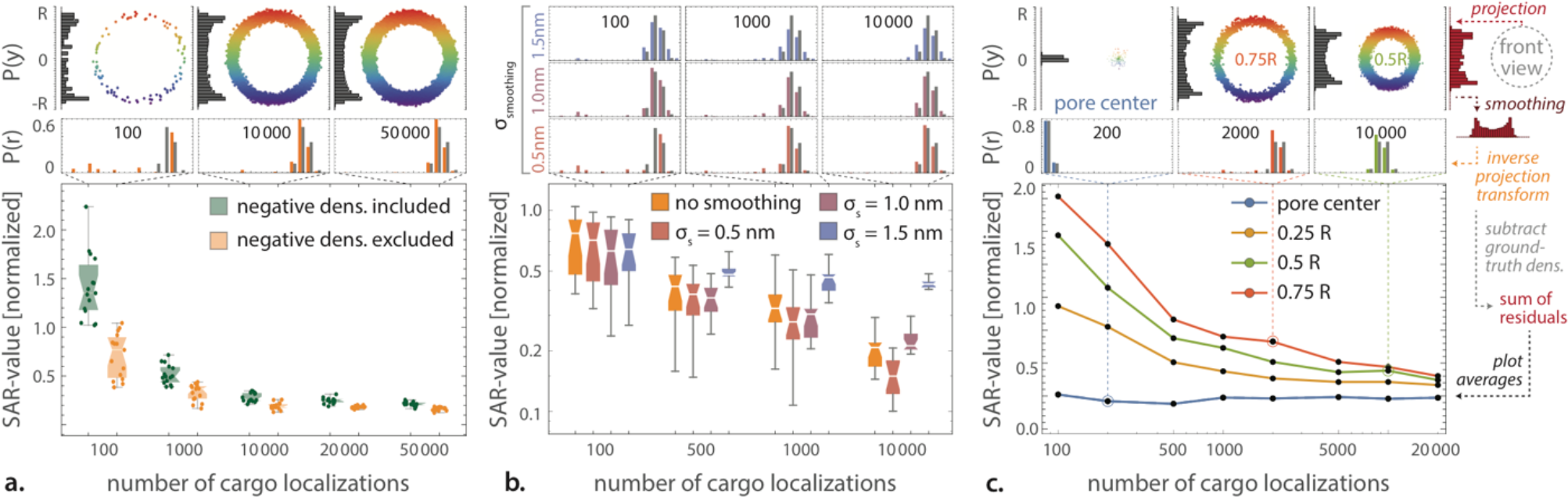
Reconstruction quality depends on dataset size, the amount of smoothing and the shape of the cargo distribution. An increasing number of cargo localizations results in a smaller and more consistent residual sum (**a**, lower chart) and hence a better reconstruction of the radial cargo density as indicated by a decreasing sum of the absolute residuals (SAR-value). Three representative simulated datasets are shown (**a**, top panels) along with their projected density profiles (grey vertical histograms) and reconstructed as well as ground truth radial density profiles (orange and gray histograms, respectively). As the number of cargo localizations in the dataset increases, the reconstruction transformation becomes more accurate, resulting in a convergence of the reconstruction error - with and without absolute negative densities. The SAR-values can also be reduced by smoothing the projected density profiles before reconstruction, often resulting in more accurate reconstructions – primarily for small datasets (**b**, lower graph; representative examples for different conditions shown in top panels). The shape of the radial cargo distribution also influences the quality of reconstruction: Gaussian cargo distributions that peak around the center of the channel are constructed with greater likeness than those that peak closer to the periphery and are less dependent on the number of cargo localizations (**c**, lower graph; three representative simulated reconstructions are shown in the top panels). Each condition in the lower three charts represents 15 independently generated datasets. This simulation assumed infinite precision (σ_tot_ = 0), perfectly aligned NPCs (orthogonal projections) and zero experimental errors.

Secondly, in order to obtain sufficient data-points to render the reconstruction of the radial cargo distribution possible, Yang et al. outline a procedure to combine cargo localizations from multiple NPCs through a data-registration (summarized in **Figure S1b-c)**. The registration process uses the centroid of the NPC marker signal to try to maximize overlap **( Figure S4a-c)**; the reconstruction quality that can be achieved therefore depends on the number of labelled nucleoporins and their distribution within the NPC to estimate the central axis. While this was paid little attention to in the original work, our simulations indicate that even with a large number of labeled nucleoporins (> 10 per NPC) a registration precision of only roughly 5 nm is achieved **(Figure S4d-e)**. This is troublesome, since our analysis shows that registration inaccuracies as small as <3nm nanometers can already produce misleading reconstruction results **(Figure S4f)**. The problem is further aggravated by the relative rotation that different NPCs may have with respect to one another: although the authors claim the ability to detect in-plane NPC rotation as small as 1.4 degrees ^1–5^, we could not reliably detect in-plane NPC rotations smaller than 20 degrees for idealized, noise-free data **(Figure S5b)**. The inability to detect in-plane rotations of a few degrees can result in high reconstruction errors as indicated by our simulations **(Figure S5c)**. Precise detection of the NPC orientation is critical to the establishment of 3D-SPEED data as ‘pseudo’-tomographic method, because the major requirement of a model-based 3D-reconstruction using single 2D-perspective, is that the orientation is known.

**Figure S4.**
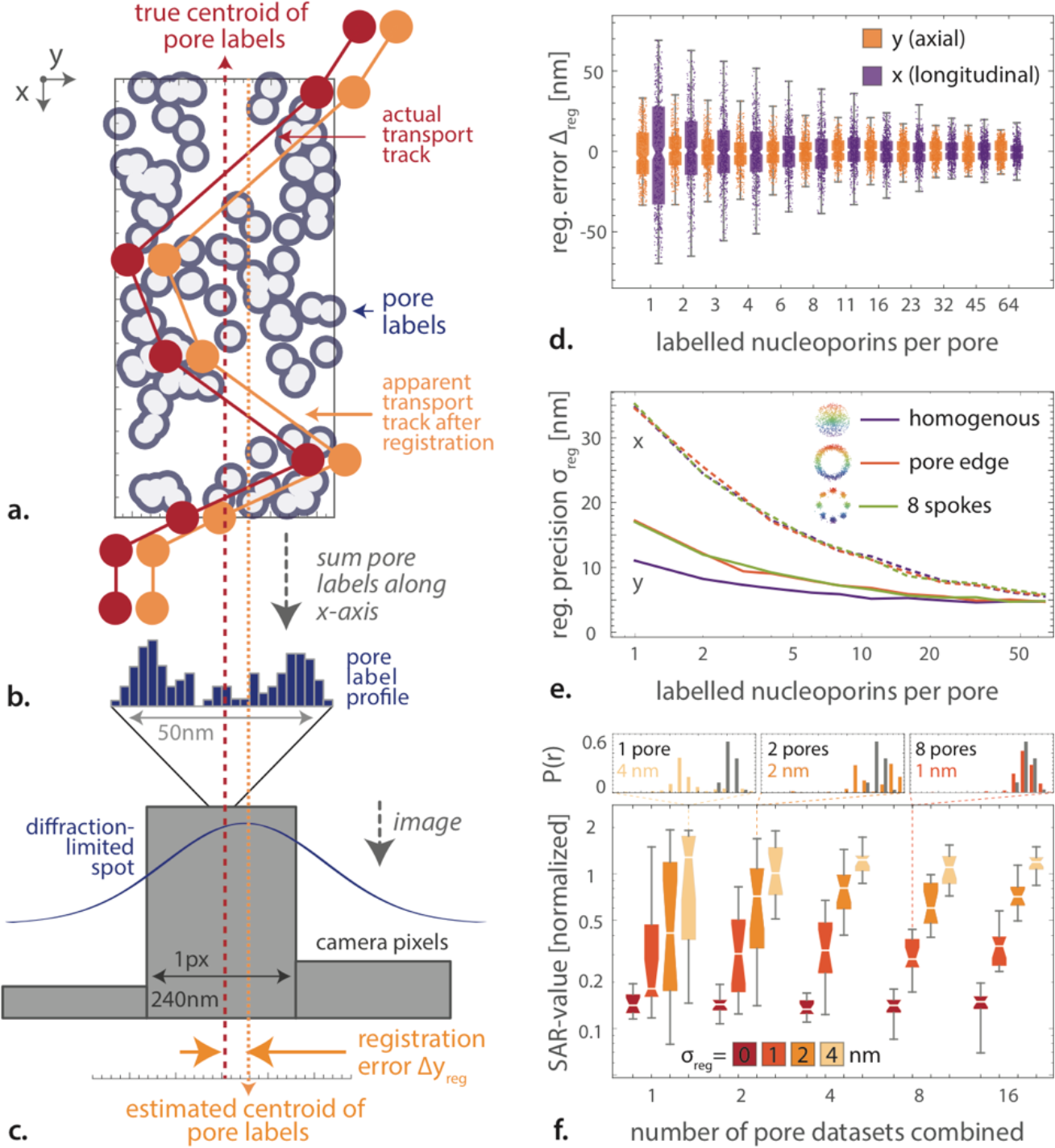
Precise registration is required for accurate cargo density reconstruction. The number and distribution of fluorescent tags in NPCs affect the pore registration precision. Fluorescent tags at C-terminus of labelled Pom121 (**a**, grey circles) can be detected as super-imposed diffraction-limited spots and subsequently used to determine the axis of the central channel (**b,c**) through either centroid extraction or Gaussian fitting. The availability of a limited number of labelled nucleoporins can result in an inaccurately estimated central axis (orange dotted line) and subsequent registration errors, causing the apparent cargo tracks (orange circles) to be shifted with respect to the actual cargo tracks (red circles). By simulating the registration error for 500 NPCs per condition (**d**), the average registration precision can be determined (e; x-registration solid, y-registration dashed). The distribution of the labelled nucleoporins inside the central channel has a slight effect on the axial localization precision, whereas the number of labelled nucleoporins has a bigger impact on both the axial and the longitudinal registration precision (**e**). When performing reconstructions on registered simulated datasets as described in **Figure S2,** we find that registration inaccuracies as low as a few nanometers can results in poor reconstruction qualities as measured using the sum of absolute-valued residuals (**f**, bottom chart). As becomes apparent in the highlighted examples (**f**, top panels), shifts due to registration inaccuracies can be clearly noted. Colors in (**f**) represents the simulated registration precision **σ**_reg_; each condition in the box-whisker was repeated for 15 independently simulated datasets, each containing 10^3^ cargo localizations.

**Figure S5.**
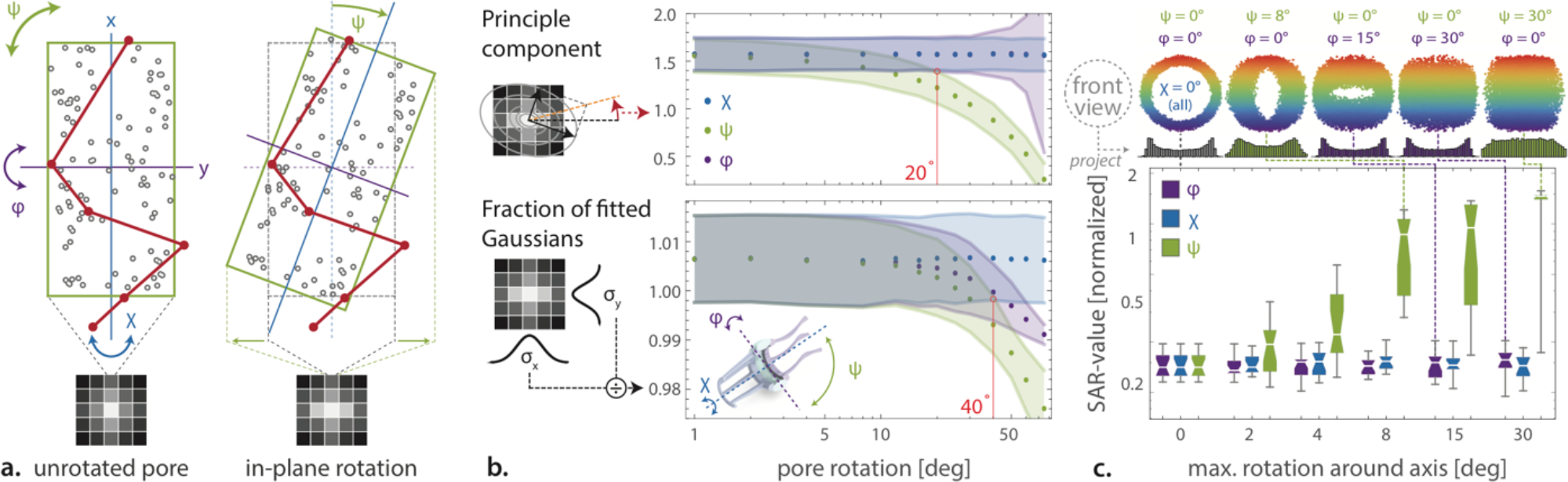
Small deviations in projection angle can greatly reduce reconstruction quality of the radial cargo density when undetected. The orientation of the NPC with respect to the observed projection can be defined as a rotation (χ,φ,ψ) around the x, y and z-axis, respectively **(a**). The rotation-estimation method presented by Yang et al. (based on determining the fraction of the widths of two fitted Gaussians, (**b**, bottom)) as well as our improved method (using principle component analysis of a fitted 2D Gaussian, (**b**, top)) are unable to reliably detect angles in the single-degree regime (**b**; 95% predication intervals shown). By performing radial density reconstructions on simulated datasets under rotation, it becomes evident that an undetected rotation of even a few degrees can severely reduce the reconstruction quality of the radial cargo distribution profile as measured using the sum of the absolute residuals (SAR-value; c, bottom graph). As expected, in-plane rotation (ψ) is the only rotational direction to affect reconstruction quality, as rotational symmetry around the pore axis (x) is assumed and only a single longitudinal zone was considered for the purpose of this simulation. The box-whisker plot in (**c**) is the result of 15 independently simulated datasets per condition; the top panels show the apparent orientation of the cargo detections (colored dots) and the resulting projection profiles (bar charts) for five representative simulations under different angles of NPC rotation.

Thirdly, the overall localization precision of cargo affects the reconstruction quality of the rotational distribution. Localization precision is a measure for the degree of certainty that can be reached when extracting the projected positions of single-molecule cargo signals from the image data and is influenced by a large number of factors including the localization precision, registration precision (between color channels as well as combined NPC measurements) and the fluorescent labelling strategy (fluorophore size, linker length etc.).

The current literature on 3D SPEED is unclear in terms of achieved localization precision as discussed by ^10^. A systematic study of the correlation between image acquisition speed and localization precision finds that for frame rates faster than 1 ms (typically 0.4 ms for 3D SPEED), the localization precision is limited to > 10 nm ^11,12^. Here we investigate the reconstruction quality using simulated cargo localization datasets with different localization precisions **(Figure S6)**. Using an error-free (i.e. 0 nm overall precision), sufficiently large (thousands of localizations) dataset, an informative and reliable reconstruction can be obtained **(Figure S6a)**. For localization precisions as reported ^1–6^, the resulting reconstructions are neither accurate nor reconstruct informative radial cargo densities **(Figure S6c)**. In fact, overall precision values of just a couple of nanometers generally result in SAR values on the order of unity or larger **(Figure S6b)**, meaning that the sum of the residuals (total error) exceeds the sum of the signal. As these simulations – even those that include simulated localization imprecision - reflect an idealized measurement system, we cannot explain how this method could produce informative results under the reported experimental conditions.

**Figure S6.**
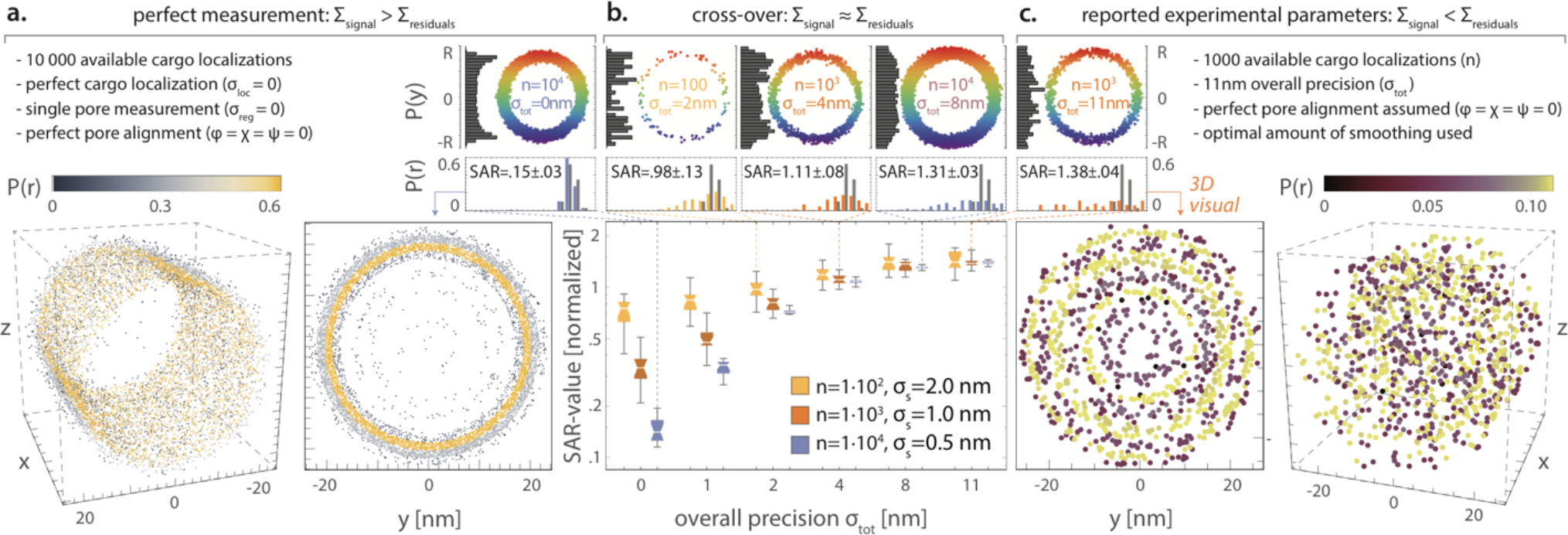
Dataset size and overall precision determine the quality of radial cargo density reconstructions. Assuming a perfectly positioned pore (i.e. perfect alignment with the pixel grid), the number of cargo localizations and the overall localization precision (a function of single molecule localization precision, measurement registration precision, drift etc.) determine reconstruction quality of the radial cargo distribution. While a reconstructed radial cargo distribution obtained under ideal conditions (top-left corner) can be informative (**a**), the reported experimental parameters (top-right corner) are not sufficient to obtain a truthful density reconstruction (**c**). Looking into the parameter-space between these regimes, various borderline-informative combinations of dataset size n and overall precision σ_tot_ can be identified (**b**, top 3 panels) and quantified using the sum of absolute residual values for 15 independent simulations (**b**, bottom graph). The visualizations in the bottom panels of (**a**) and (**c**) show data-points randomly generated using the reconstructed radial cargo distributions and feature the same number of particles as the initial datasets (small top panels; rainbow colored). Bar-charts in the five reconstruction panels are presented as in **Figure S2**; SAR-value insets represent mean±SD for 15 independent simulations.

Fourthly, model-assumptions such as cylindrical symmetry can lead to misleading reconstructed cargo distributions if they are not fully met. The 3D SPEED method does not predict a particular 3D-structure but rather projects data onto a solution space that is limited to cylindrically symmetrical solutions. While it is possible that cargo particles occupy the nuclear pore in cylindrically symmetrical distributions, this is by no means the only possibility. If nuclear the fibrils would homogenously occupy the central channel and no other interactions would exist, the resulting homogeneity could result in cylindrical symmetry of the cargo distribution. However, an eight-fold symmetry of the NPC is generally accepted ^13–15^. If transporters would follow this symmetry, the transport distribution would exhibit an eight-fold point-symmetry that approaches cylindrical symmetry, but only to a limited extent. Notably, both dynamical and inhomogeneous FG meshwork as well as different point-symmetries (i.e. numbers of spokes) have been reported.^16–20^ Another potential source of cylindrical symmetry was suggested in the form of super-imposed datasets of NPCs that freely rotate around the transport axis, which would average out asymmetries. While possible in theory, free rotation of NPCs has not been demonstrated and this workaround would require accurate superposition of a large number of measurements.

To assess the potential implications of studying non-symmetrical cargo distributions using this method, we performed reconstructions on simulated NPC cargo localization datasets that exhibit varying amounts of cylindrical symmetry **(Figure S7)**. Our simulations show that for symmetry violations - even at the level of preferred travel of cargo along 6-12 ‘spokes’ - the reconstructed cargo distributions can be both inaccurate and misleading to a degree that depends strongly on the imaging perspective (i.e. the relative rotation of the NPC – see **Figure S5b)**.

**Figure S7.**
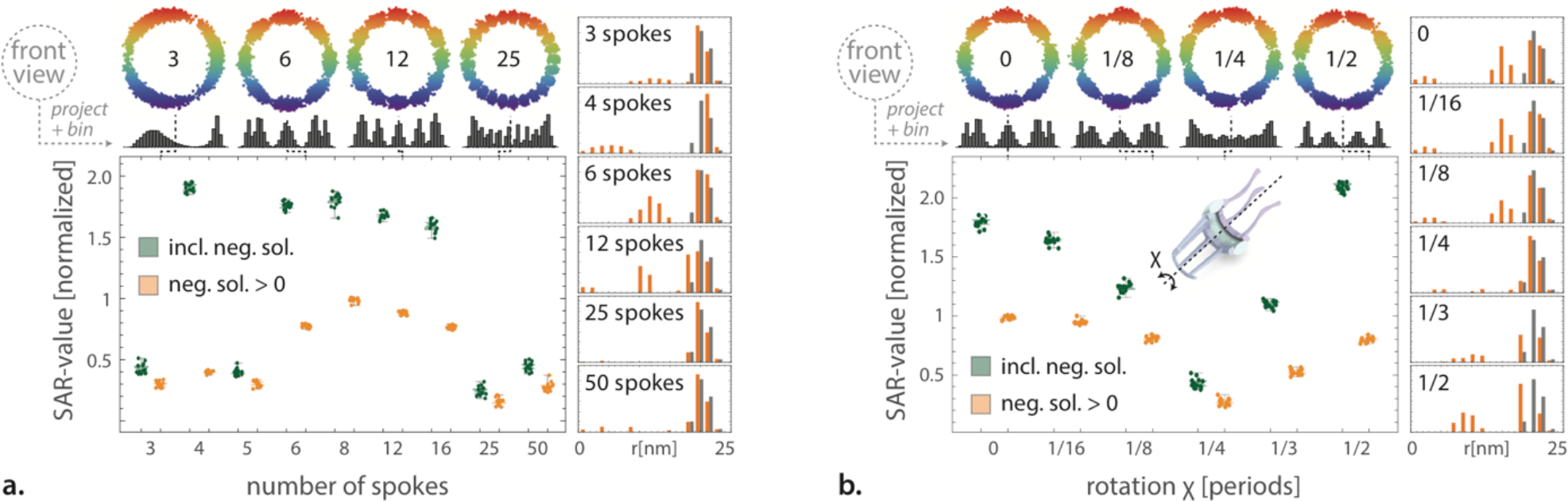
The assumption of cylindrically symmetrical cargo distributions can yield result in misleading reconstructions. If one is to reconstruct the radial cargo density from localizations in a single projection, the underlying distribution needs to be cylindrically symmetric. By simulating cargo localizations with varying degrees of cylindrical symmetry, significant sensitivity is found for strong violation of the symmetry assumption (low number of spokes) as measured by the sum of absolute-valued residuals (SAR-value) (**a**; bottom-left chart). Representative examples of the simulated cargo localizations (top panels, colored dots), the corresponding binned projections (top panels, grey histograms) and the resulting reconstructions (right panels, orange bars) show how applying the reconstruction process to asymmetric distributions can produce misleading solutions. A similar conclusion can be drawn when varying the angle of projection in case of an asymmetrical cargo distribution (**b**; rotation **χ** around the central axis of the pore is denoted as the fraction of a single period for an eight-fold symmetry). Each condition was repeated 15 times using independently simulated datasets of 10^18^ cargo localizations each; no errors or inaccuracies were simulated.

In summary, we have shown that the 3D SPEED method would work for sufficiently symmetrical distributions, given that the dataset is large enough (thousands of localizations) and of sufficient overall precision is achieved (a few nanometers at most – see Figure S6b). Starting with the assumption of perfect data onto which we introduce known, small errors, we show that it is highly unlikely that any 3D-SPEED data as recorded under reported conditions would be able to produce accurate cargo distribution reconstructions. To come to this conclusion, we ignored practical challenges that have been disputed in the past, such as the questionable ability to image a single NPCs ^19,21^, or the presence of chromatic aberrations ^16,17,19,21^. Furthermore, the use of a tilted illumination is likely to result in PSF deformations, which would need to be accounted for in order to reach an acceptable localization precision and eliminate any biases. Similarly, it has been shown that heavy spatial under-sampling of the fluorescent signals on the camera, as is the case in the experimental data published, can result in a localization bias of tens to hundreds of nanometers ^22^. Additionally, it should be noted that we considered the NPC as a single longitudinal segment throughout our evaluation. Yang and colleagues have repeatedly split the cargo localization dataset into multiple segments and performed individual reconstructions on each of them (illustrated in Figure S1c-f). If done judiciously, differences cargo distributions could be examined at different spatial steps of the transport process, increasing the amount of information that can be obtained from a single dataset. While theoretically possible, such treatment would strongly increase the number of data points required for reconstruction and would place even more stringent requirements on the rotation detection and registration precision – none of which are currently met. Another consideration here pertains to the sampling of such multiple distributions: transport molecules tend to spend more time at the NPC in some states than others (e.g. docking vs translocation), affecting spatial sampling accordingly ^23^. The use of probability-based detection methods - rather than the 2D-Gaussian fitting used here - would increase the detection efficiency and help increase confidence in the quality of the dataset by allowing control of the false-positive rate^11,12^. As a final remark, we note that a spatial mapping of 2D image data is not a tomography method as suggested by the creators of 3D-SPEED^4^- neither by definition, nor by resolvable data content. The proposed method does not extract 3D information; it merely facilitates a coordinate transformation based on model assumptions.

### Mathematical description of the model-based reconstruction

This chapter provides a mathematical description of the proposed model-based reconstruction that uses projected transport localizations in order to reconstruct the radial distribution of molecules within the central channel of the nuclear pore complex (NPC).

#### The Reconstruction Equation

We adopt the model proposed in the original 2010 publication^1^ which assumes that a NPC exhibits a circular cross section and can be modeled as a cylinder with the local coordinates defined as shown in **Figure S8a**. To define the equation for reconstructing the 3D density distribution, we assume that the underlying probability density is a function of the radial position within the pore and independent of angle with respect to the axis of the central channel and constant along the length of the pore segment (cylindrical symmetry). Then we embed the NPC into a rectangular box **(Figure S8b)**. In the XY-plane, the projection image can be divided into N parallel sections along the X-axis. The α-^th^ section is denoted by *S*_α_ and has length |*S_α_*|. Each segment *S*_α_ can now be partitioned in two ways: using rectangular or circular partitions (I-partition and R-partition in short **(Figure S8b))**. For the fixed section *S*_α_, the rectangular region is divided into M parallel strips {*I_1_,I_2_,*…I_*M*+1_} with boundaries {*a*_0_, *a*_1_,… *a*_*M*+1_} and *a_0_ < a_1_ <*…< *a_M+1_* while the cross section is divided by annuli {*R_1_,R_2_,…R_M_*} with radii {*r_1_,r_2_,…r_M_*} where *r_1_ < r_2_ <* < *r_M_ = R*, *R* is the radius of the NPC cylinder and *R_M+_*_1_ denotes the region outside the NPC cylinder. The intersection region of the strip *I_i_* and the annulus *R_j_* is denoted by *A_ij_* **(Figure S9a)**. For each section *S_α_*, *w_k_* and *f_m_* are defined to be the number of data points in *I_k_* and the density of data points in the annulus *R_m_*, respectively. *f_m_* is hence assumed to be constant in the annulus *R_m_* for *m = 1,…,M* and *R_M+_*_1_.

**Figure S8.**
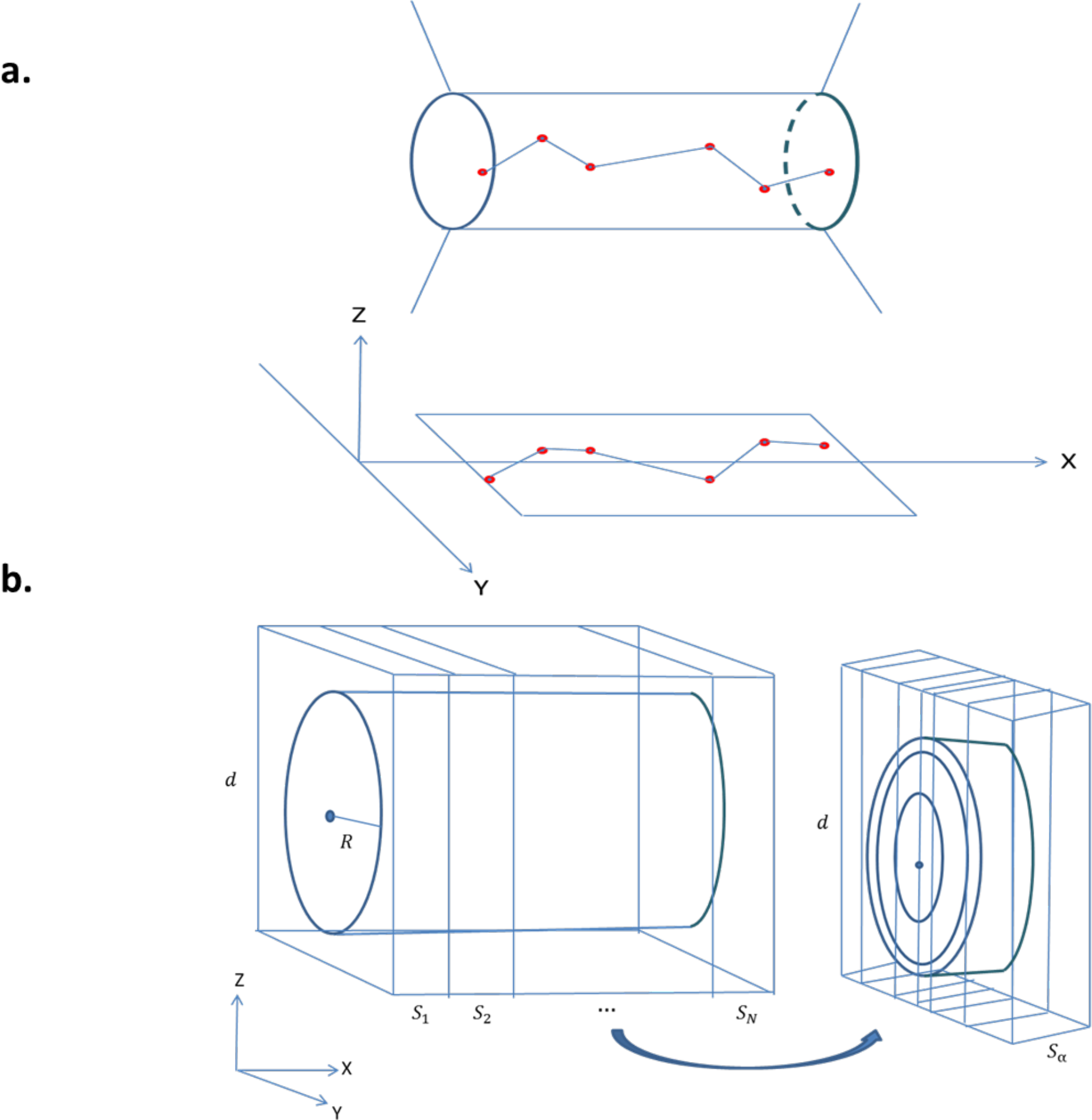
Setup of the geometry of NPC, projection plane and partitions. (**a**) Schematic diagram of the 3D cylindrical NPC,2D projection plane and Cartesian coordinate system. The NPC is modeled as a cylinder with circular cross section. The red dots represent cargo localizations in 3D and in 2D projection along Z-axis during cargos transport through the NPC.(**b**) The partition of NPC. Left: The cylindrical NPC is embedded into a rectangular box where d is the length of the box along Z-axis and R is the radius of the NPC. Along X-axis the box is divided by N sections, denoted by S_1_…,S_N_. Right: For each section, say Sα, the box is partitioned along Y-axis, called I-partition and the contained NPC cylinder is divided by annuli, named R-partition.

**Figure S9.**
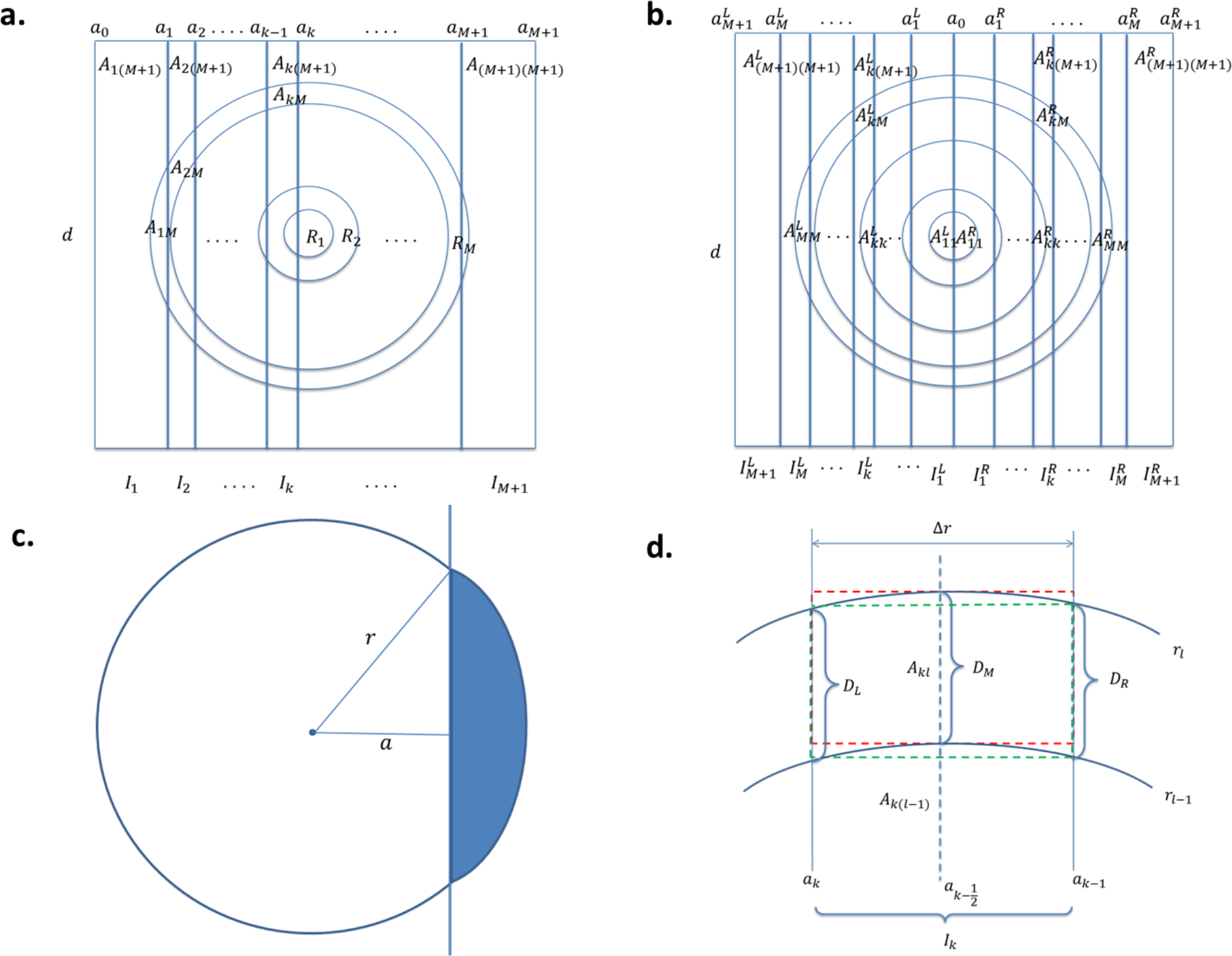
I- and R-Partitions and associated areas. **(a)** General partitions: the rectangular cross section of NPC box is divided into M+1 parallel strips {I_1_,I_2_,**…** I_m+1_} with boundaries {a_0_, a**1**,… a_M+1_} while the circular cross section of the NPC cylinder is divided by annuli {R_1_, R_2_,**…** R_M_}. The region outside the NPC cylinder is denoted by R_M+1_. A_ij_ is defined as the intersection region of the strip I_i_ and the annulus R_j_. (**b**) Symmetric partition: The partition is symmetric with respect to the central line a_0_. The superscripts L and R denote the quantities on the left and right sides. (**c**) Circular Segment: The area of the shaded circular segment is denoted by S(r,a) and computed by r^2^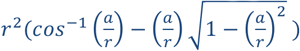 The area of each A_ij_ can be computed by the alternative sum of S(r,a) for some r and a. (**d**) Area of A_kl_ and first-order approximations: The exact area of the region A_kl_ bounded by two lines a_k_ and a_k-1_ and two circles with radius r_l_ and r_l−1_ can be computed by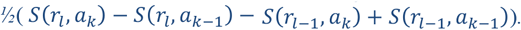. The rectangles marked by green and red dash lines are first-order approximations of A_kl_ and the areas are computed by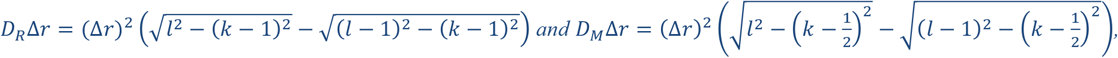 respectively.

By using the relation that the number of data points in a space is equal to the product of the density of data points and the volume of the space, we may obtain the following matrix equation for the section S_α_ ^1^,

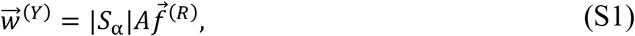

where 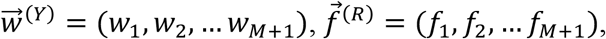, and the entry of the matrix *A* is given by the area of *A_ij_* which encodes the information of the partitions. Thus, we call *A* the partition matrix. To solve the Eq. (S1), it is required that matrix *A* be invertible. The invertibility of the matrix *A* is determined by the partitions. In what follows, we omit the factor |*S*_α_| to simplify the notations and consider a special kind of partitions, symmetric partitions (Figure S9b), which are slightly more general than the equipartition used in the original work^1^ and can be shown to produce invertible partition matrices.

In a symmetric partition, it is required that the all strips are symmetric with respect to the central line *a*_0_ which passes through the center of the circular cross section **(Figure S9b)**. This symmetry implies that 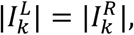, and 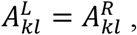, for all *k,l* = 1,2,…, *M* + 1 where the notation |*I*| denotes the width of the strip *I*. However, in practice the set of data points may not develop exactly radial symmetry (or any symmetry) and then 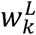 in 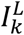 and 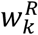 in 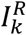are generally not equal. To mitigate this inconsistency, we may sum 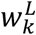 and 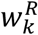 together and let *A_kl_ =* 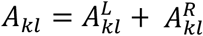 for all *k* and *l*. By doing this, we may write

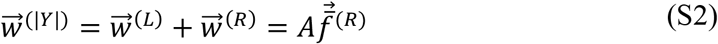

Where 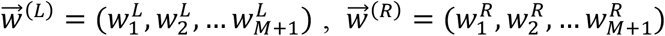 and 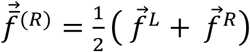 is the average density vector of the left and right density vectors. In addition, we impose the constraint 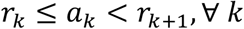 on the partitions. With this constraint, the partition matrix becomes an upper triangular matrix as follows:

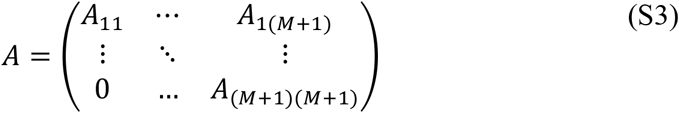

where by abuse of notation, we denote the entry of the partition matrix *A_ij_* the area of the intersection region *A_ij_*. This upper triangular matrix is invertible. The equipartition used in [1] is a special case of the symmetric partition, namely, 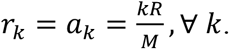.

With given I- and R-partition, we may calculate the area of *A_ij_.* It is useful to define *S*(*r, a*) be the area of the shaded circular segment as shown in **Figure S9c** and compute

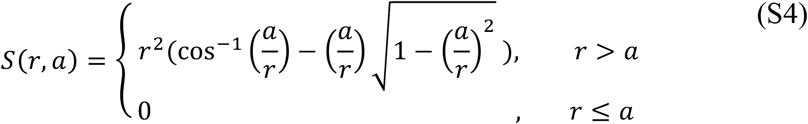

The area of *A_ij_* can be expressed by the alternating sum of the area of appropriate circular segments. In fact, the entry of the partition matrix *A* is given by

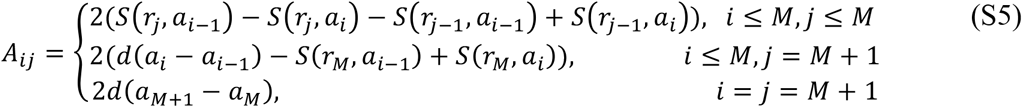

The area formula in Eq. (S5) is slightly different from that used in the original work^1^. In fact, the expression in Eq. (S5) is the exact area of *A_ij_* while the area formula used in the original publication^1^ is a first order approximation of the Eq. (S5). To see that, let us consider the following first order approximation by Taylor expansion in a small parameter ϵ,

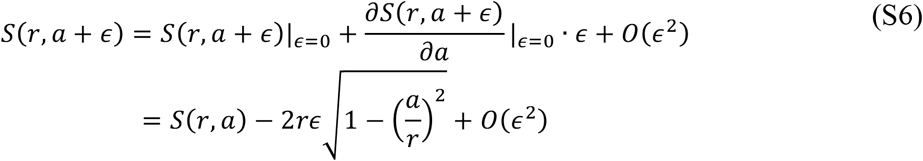

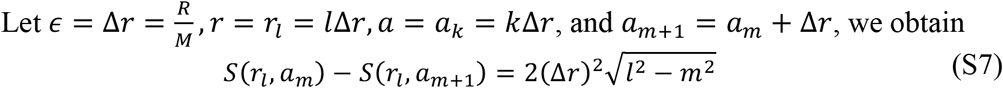

Therefore,

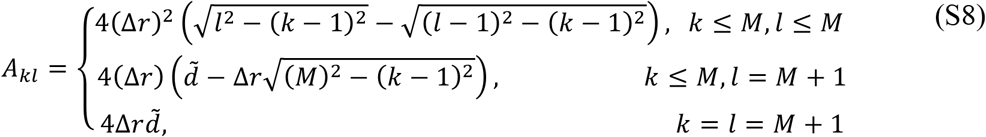

where 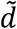 is defined as the same as *d* in the original work^1^ and 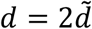 has been used. The factor 4 comes from the fact that in the original work the area formula is applied to one quarter of the cross section while we consider the full cross section.

There are several options for the first order approximations of the Eq. (S5) **(Figure S9d)**. The Eq. (S8) describes the area of 4*D_R_Δr* in **Figure S9d** and the area formula used in the original work^1^ is given by

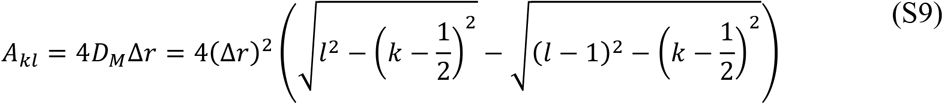

Alternatively, 4*D_L_*Δ*r* and 2(*D_R_* + *D_L_*)Δ*r* can be options to approximate the area of *A_kl_*. The accuracy of these approximations depends on the number of the partitions *M*. The lager *M* is, the better the approximation.

#### Deformation of Partitions and the Space of Solutions

By inspecting the Eq. (S1), 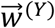 is complete determined by I-partition. When the I-partition is deformed through changing the position of boundaries {*a*_1_, *a*_2_,̤ *a*_M+1_},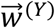 and the entry of the partition matrix *A* are changed as wells as the density vector 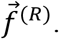. In general, three terms in the Eq. (S1) are varied simultaneously by this I-deformation. while keeping 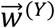 untouched, which makes it easier to investigate the structure of the space of solutions. That is the R-deformation, namely deforming the R-partition by varying the radius of annuli. Here we illustrate the main idea by using a simple example, which can be generalized for more complex configurations. Let us consider a 2-partiton reconstruction equation where *A* is a 2x2 matrix, and the Eq. (S1) takes the form

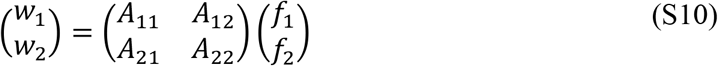

Before applying R-deformations, we fix an I-partition and then *D*_1_: = *A*_11_ + *A*_12_ and *D*_2_ **≔** *A*_21_ + *A*_22_ are fixed where *D*_1_ and *D*_2_ are the area of the strips *I*_1_ and *I*_2_ in the I-partition, respectively. When the R-partition is deformed, the coefficients *A*_11_, *A*_12_, *A*_21_ and *A*_22_ are varied but subject to the constraints *A*_11_+*A*_12_ = *D*_1_ and *A*_21_+*A*_22_ = *D*_2_. Geometrically, this deformation changes the slope of the lines *A*_11_*f*_1_ + *A*_12_*f*_2_ = *w*_1_ and *A*_21_*f*_1_ + *A*_22_*f*_2_ =*w*_2_. As looking into the Eq. (S10) carefully, we notice that this deformation has two fixed points, 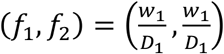 and 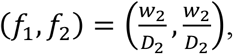, and that the R-deformation in this case rotates the lines around their respective fixed points. Notice that the positions of fixed points are determined by the I-deformation. The space of the solutions is then the intersection of the deformation region of the lines **(Figure S10)** where we introduce two deformation parameters *t* and *s* defined by *t* = *A*_11_ and *s* = *A*_21_, respectively. In general, for a 2x2 matrix *A,* the space of the positive solutions forms a polygon **(Figure S10)**. When *A* is an upper triangular matrix, the solution space reduces to a one-dimensional segment. For higher rank cases, the solution spaces should form polyhedrons. From **Figure S10**, we can see that typically the positive solutions of the Eq. (S1) with given 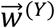 are not isolated points in the *f*_1_*f*_2_-plane; instead they form a space with nonzero dimensions. More importantly, these solutions are mathematically valid reconstructed densities, unless more principles or selection rules are imposed. That is to say, the reconstructed densities 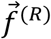 are not unique but infinite. Besides, it also can be seen from **Figure S10** that there are two disconnected branches of solution space: in each branch the negative solutions are much more ample than positive ones and these two sets of solutions are connected. With the connectedness, positive solutions can be found by continuously deforming the R-partition along an appropriate line, such as *L*_1_ or *L*_2_ whenever a negative solution is solved in the initial partitions.

**Figure S10.**
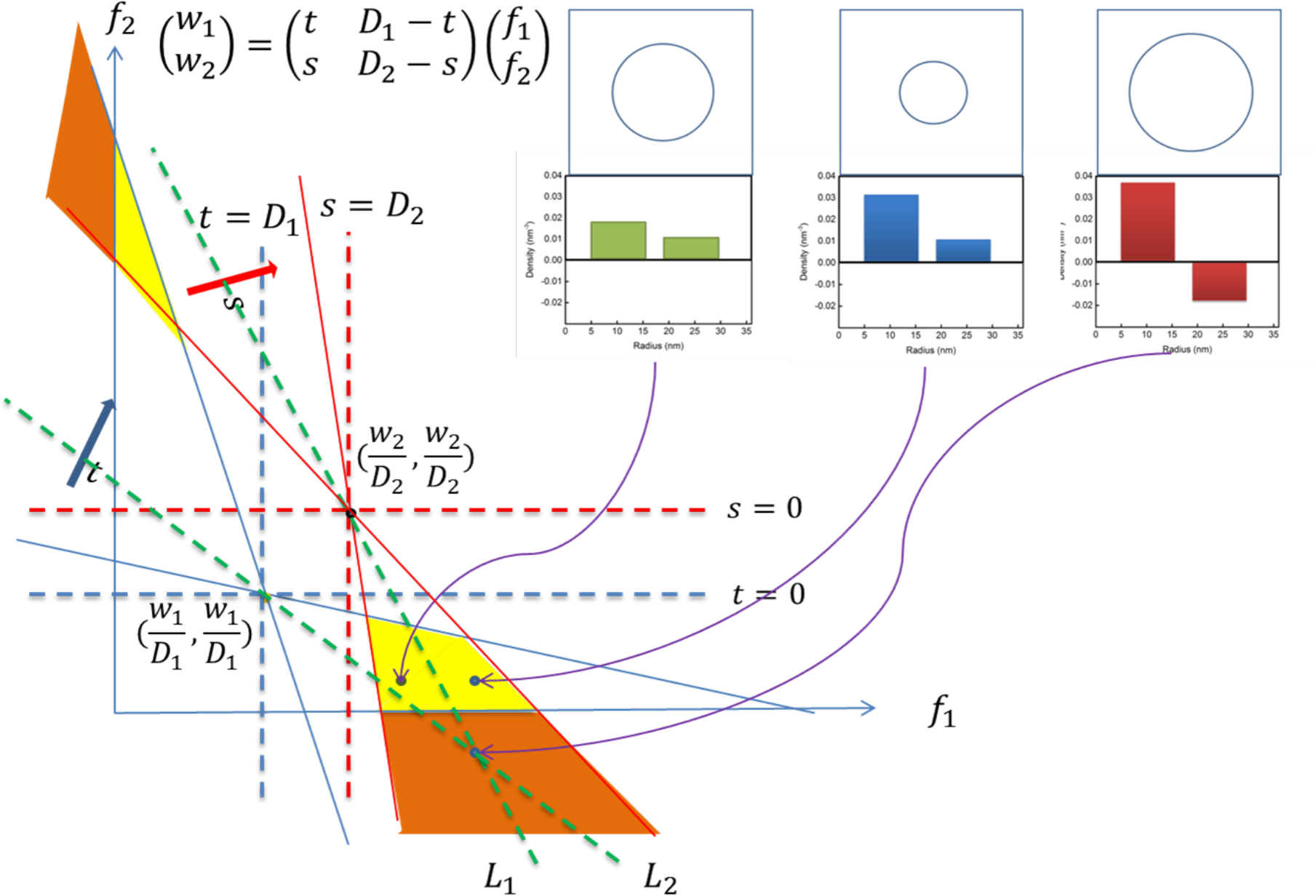
The solution space of 2-partition reconstruction equation and R-deformation. The 2-partition reconstruction equation is a system of two linear equations. To demonstrate the R-deformation, parameters s and t are introduced while I-partition are fixed, since D_1_ and D_2_ are constants. The intersection of two red dash lines and of two blue dash lines determine two fixed points of the R-deformation parameterized by s and t. When the parameter s (t) varies, the red (blue) solid line is deformed by rotating red (blue) solid line with respect to the fixed point as indicated by the red (blue) arrow. The solution space is determined by the overlap of s- and t-deformations. In particular, the yellow and orange regions indicate the solution space of positive (f_1_, f_2_>0) and negative solutions (f_1_ or f_2_<0), respectively. A negative solution in the orange region (for example, the point in the intersection of green dash lines L_1_ and L_2_) can be deformed into a position solution in the yellow region by rotating L_1_ or L_2_ with respect to the individual fixed points.

#### Transformation of the Normalized Distributions P(|y|) and P(r)

With the Eq. (S1), we can drive the associated equation relating normalized distributions *P*(**|***y***|**) and *P(r)* defined on the Y-axis and radial direction, respectively. Let us define the normalized distribution on the Y-axis by 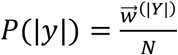 where ***N*** is the total number of the cargo localizations. Then

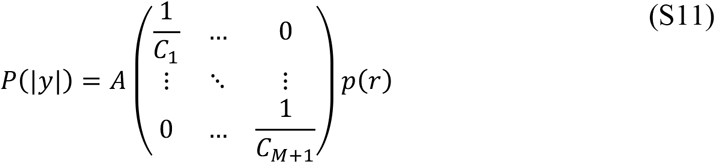

where *C_k_* is the area of *R_k_*. Therefore,

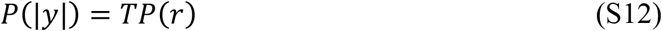

where

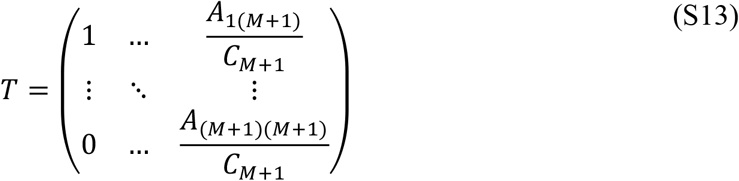

By the relation, 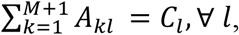, we found that summing up all components in each column vector of the matrix *T* is equal to one. This matrix *T* can be treated as a probability matrix and is determined completely by the partitions. From the Eq. (S12), we can see that the normalized distribution *P* (|*y*|) on the Y-axis is not the same as the normalized distribution *P(r*) on the radial direction. Instead they are related by the probability matrix *T* which can be varied by deformations. In particular, *A* and *P(r)* on the right-hand side of the Eq. (S12) can be deformed while *P(*|*y*|) remains invariant under the R-deformation. This makes sense, since the 2D image data is a physical quantity, which can be measured and thus be regarded as the average of the density profiles on the radial direction. This conclusion also indicates that the normalized reconstructed distribution *P(r)* is not unique for given 2D image data. This is the same conclusion arrived at through the study of the space of solutions in the previous section.

In conclusion, we have demonstrated that back-projection transformation in 3D SPEED produces non-unique reconstructions that require careful conditioning to produce meaningful, positive and artefact-free distributions. Even if performed in a mathematically sound way, there are numerous error-contributing factors such as undersampling and localization imprecision that make truthful reconstructions using 3D SPEED challenging, if not impossible under state-of-the-art measurement conditions.

### Simulation Methods

Simulations were performed using Wolfram Mathematica 10 and 11. The annotated source code notebooks will be made available upon request.

The simulations can be divided into two main groups: those looking at reconstructed *densities* and those looking at reconstructed *distributions* of transported particles. While the former was used to allow direct comparison with published work (see **Figure 1)**, the latter is more suitable for benchmarking and optimization as it can be readily compared to the input distribution of the simulation (all supplemental figures). The simulation implementations are quite similar for both approaches, but there are some differences in the simulation parameters and data interpretation which are outlined in the final sections of this chapter.

#### Radial cargo density and distribution reconstruction

To investigate the ability to reconstruct the discrete ground truth radial cargo distribution *P*_r_[*r*], we simulate *n* cargo localizations (*r*, *θ*, *x*)_n_ within the central channel as constrained by the simulated central channel volume (i.e. 0 ≤ *r* < *R*, 0 ≤ *θ* < 2π and -*l*_pore_/2 ≤ *x* ≤ *l_pore_*/2) using radial Probability Density Function (PDF) *p*_*r*_(*r*), angular distribution function *p*_θ_*(θ)* (rotation around the central channel axis) and longitudinal distribution function *p*_x_(*x*) (along the pore length). The PDFs used for the reconstruction results in each figure are listed in **Table 1**. The discrete ground truth radial cargo distribution is defined as

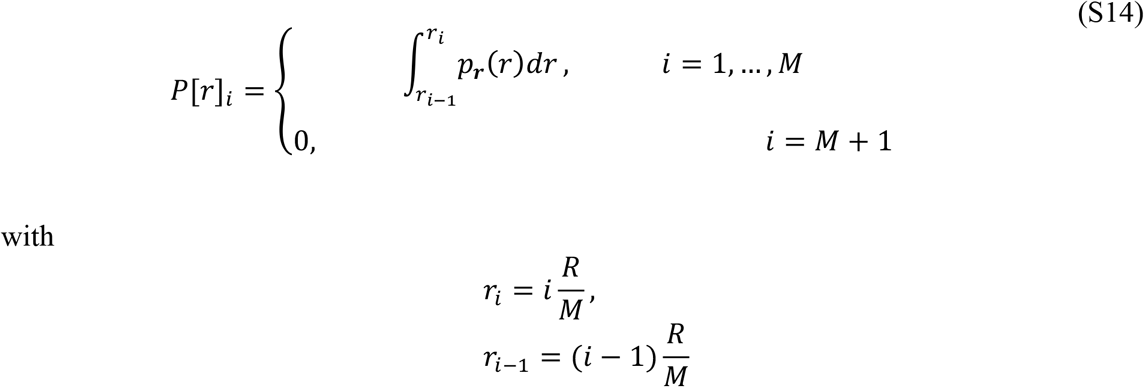

**Table 1.**
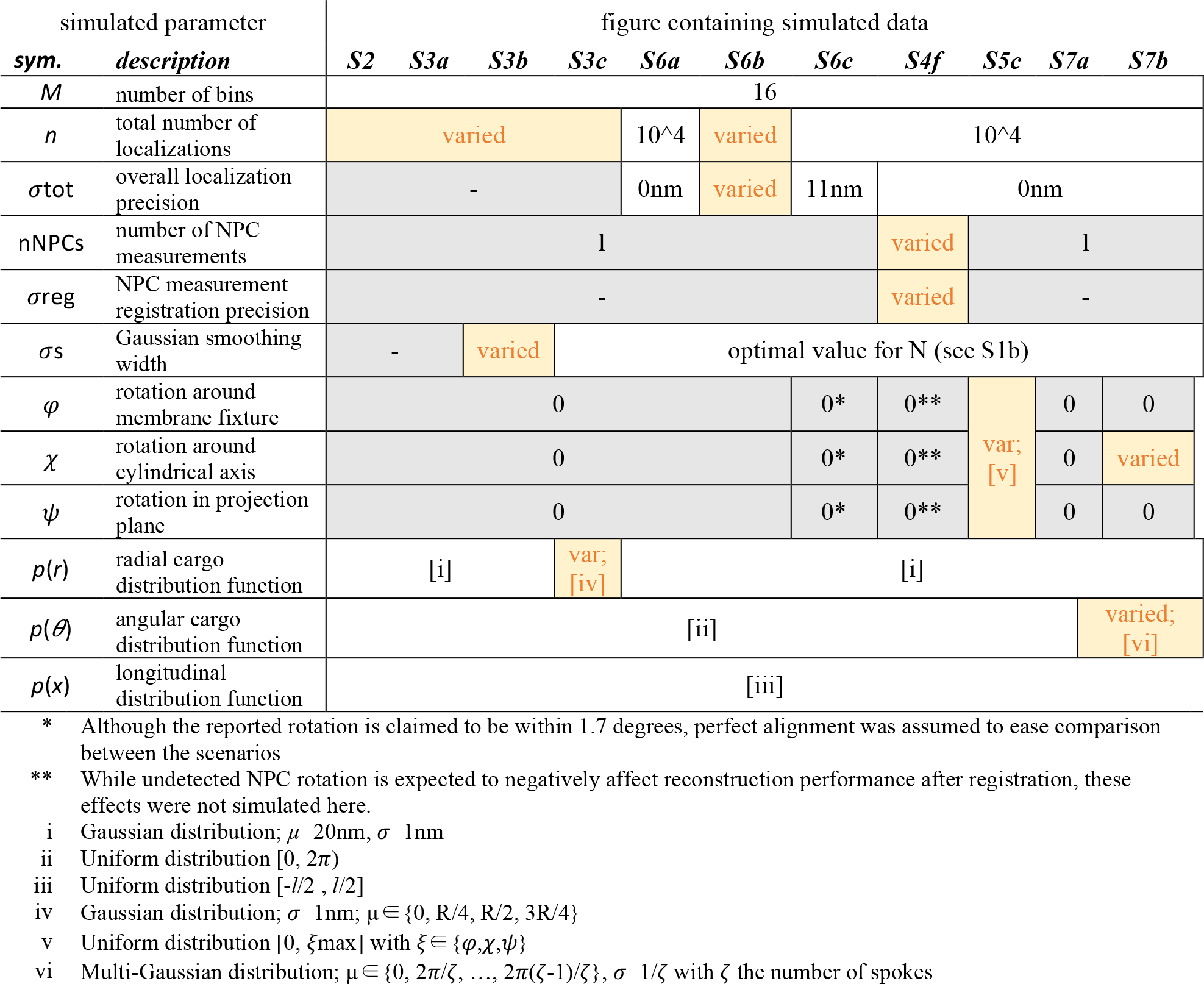
Simulation parameters for distribution-based simulations.

Cylindrical coordinates (*r*, *θ*, *x*)_n_ are then transformed to Cartesian coordinates *(x, y, z)*_n_ and subsequently subjected to a rotation transformation to simulate rotation of the entire NPC by *(χ,φ,ψ)* degrees around each Cartesian axis (X,Y,Z). To simulate localization imprecision, an offset is applied to each coordinate on a point-by-point basis to yield (*x’*, *y’, z’)*_n_ using a normal distribution with width *σ_x_=σ_y_=σ_z_ =σ_tot_.* The overall localization precision *σ_tot_* is varied between simulations and models the combined impact of localization imprecision (i.e. the precision in super-resolving the particle location through fitting), color registration, drift and other factors. The overall localization imprecision is modelled isotopically as the imaging method used in 3D-SPEED microscopy is not fully characterized in each direction; the simulated cargo localizations therefore likely represent a better localization precision than would be obtained through 3D-SPEED measurements.

The cargo localizations (*x*’, *y*’, *z*’)_*n*_ are projected onto the a single object plane disregarding their individual *z*-positions, assuming their limited range in z-coordinates (Δ*z*_*max*_ ≈ 2(*R* + 2*σ*_*tot*_) *≈* 90nm) are negligible compared to the depth-of-focus (DOF=*λ·*/NA^16^ + *n·d*_*pix*_/(M·NA) ≈ 653nm, with *λ* the wavelength, NA the numerical aperture, *n* the refractive index, *d*_pix_ the pixel size and *M* the magnification). The projected cargo localizations (*x*’, *y*’)_*n*_ are binned into 2(*M*+1) equally sized bins to yield the measured projected cargo profile *P*_*y*_’[*y*] with *y* ϵ [-(1+1/M)R,…,*-R/M, R*/*M*,…, (1+1/*M*)*R*]. Assuming cylindrical symmetry, the two halves (y<0 and y>0) may be mirror-summed to yield *P*_*|y|*_’[*y*] with *y* ϵ [*R/M, 2R/M,…*, (1+1/M)*R*]. Gaussian smoothing of width *σ*_*s*_ is applied to *P*_*|y|*_’[*y*] to yield *P*_*|y|*_”[*y*] in order to reduce the occurrence of artifacts after the Inverse Projection Transformation (IPT). The IPT is achieved by a matrix multiplication using the inverse of partition matrix *A.*

For the general case,

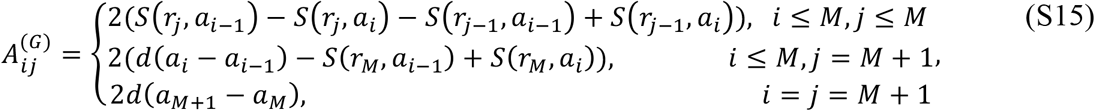

where

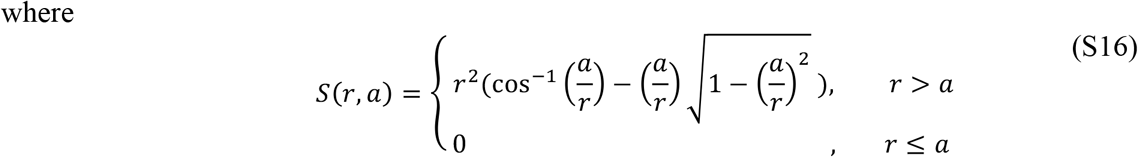

For equipartition cases, the matrix is reduced to

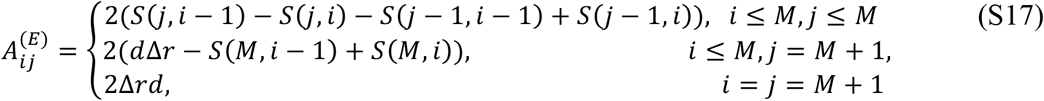

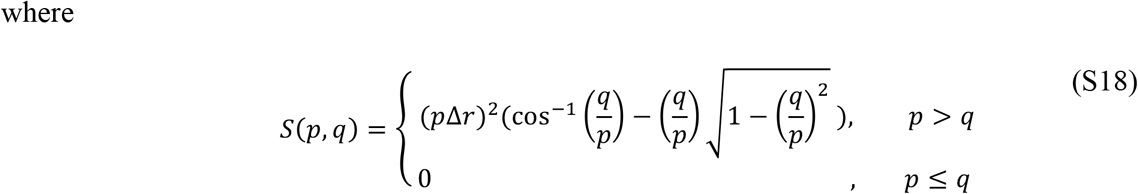

p and q are positive integers and the first order approximation used in the original work ^1^

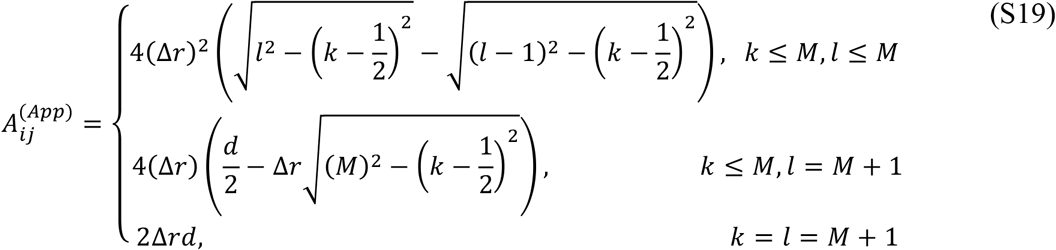

Note that 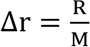 for all matrices used in simulation.

Although a range of the matrices defined above was tried for reconstruction, all results shown in this publication were obtained using 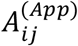 to ease comparison with published results. The inverted partition matrix was used to transform the smoothened projected cargo profile *P*_|y|_”[*y*] to the reconstructed radial cargo density *D*_r_’[*r*] using (S1); i.e.

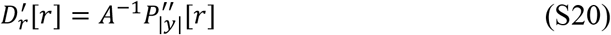

with *r* ϵ [*R*/*M*, *2R/M*,…, (1+1/*M*)*R*]. By multiplying each element of *D*_r_’[*r*] by its corresponding annular surface area and normalizing the result, the reconstructed normalized radial cargo distribution profile *P*_r_’[*r*] is obtained. The comparison of the input and reconstructed distributions rather than densities allows for a more straightforward interpretation of any discrepancies, as the number of localizations available per bin (rather than per volume) tends to have an effect on the reconstruction quality. The radial transport density, however, is potentially more useful to derive biological insights from the results; for that reason (and to facilitate comparison with published work), the simulations in **Figure 1** use radial transport densities for fitting, whereas all figures contained in the supplement present radial transport distributions.

#### Calculating the reconstruction error (SAR-value)

Despite the damping effect of smoothening the projected cargo distribution, varying degrees of artefactual negative values would result from the IPT – especially for small *n*. To quantify the truthfulness of the reconstruction independent of these artefactual densities, *P*_r_’[*r*]_i_=0 was substituted for negative values (*P*_r_’[*r*]_i_ < 0) in the reconstruction, after which *P*_r_’[*r*] was normalized again. The reconstruction error was subsequently calculated as the Sum of the Absolute-valued Residuals *(SAR)* between the discrete, normalized ground-truth radial distribution *P*_r_[*r*] and the discrete, normalized reconstructed radial distribution *P*_r_’[*r*]; i.e.

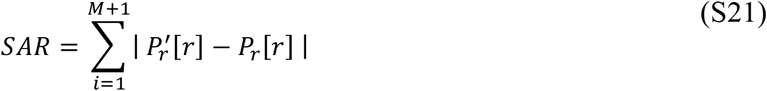

and as *P*_r_’[*r*] and *P*_r_*[r]* are both normalized, *SAR* is bound by 0 ≤ *SAR* ≤ 2.

Apart from disrupting factors arising from pore rotation and those discounted in *σ*_*tot*_, no sources of error (e.g. false-positive detections, noise, localization biases due to aberrations, drift) were simulated for; the results of these simulations are therefore likely to be over-optimistic and should be interpreted as best-case scenario outcomes for the given parameters.

#### Distribution-based-comparison simulations (suppl. Figures)

The simulation results in the supplementary figures used reconstructed particle distributions rather than densities (such as in **Figure 1)** to better facilitate quantitative evaluation of the reconstruction performance by means of the *SAR*-value. As the quality of the reconstruction tends to vary between randomly generated datasets (especially if a low number of cargo detections is available), all simulations were repeated 15 times per condition with new, randomly generated data to characterize the range of *SAR*-values resulting from a particular set of parameters. In cases where the distribution is not visualized using a scatter-plot or box-whisker diagram, the mean *SAR* value (± standard deviation) is shown.

#### Reconstructions using localizations from multiple NPC measurements

With the exception of **Figure S4f**, all cargo detections were assumed to either be obtained from a single nuclear-pore measurement or perfectly registered (*σ*_*reg*_ = 0nm) between measurements of identically oriented NPCs (*φ*_1_=φ_2_; *χ*_1_=*χ*_2_; *ψ*_1_=ψ_2_). For the multi-pore simulation in **Figure S4f,** *n*_*pores*_ datasets of *n/n*_*pores*_ localizations were simulated (*φ*=*χ*=*ψ*=0) and combined after applying a normally distributed offset of width *σ*_*x*_ =*σ*_*y*_ = *σ*_*z*_ = *σ*_*reg*_ to each individual dataset. The combined datasets were subsequently processed using the same method as described in the previous section.

#### Multi-pore registration precision

To determine the bias and precision with which the pore center is estimated (which determines the registration precision), we performed a number of simulations that explore a centroid-based method as well as 1D and 2D Gaussian fits. The properties of the simulated imaging system were taken from 3D-SPEED publications ^1–5,21,24^ and include a 100x, 1.4NA objective and a camera with 24μm pixels. The simulated wavelength corresponds to the GFP emission peak (*λ*=509nm). Simulated pores were assumed to be perfectly aligned with respect to the projected image plane and the pixel grid *(φ=χ=ψ=0)* and the simulated imaging system was assumed to be diffraction limited and noise-free.

A varying number of pore label positions was simulated according to three different radial probability density functions in the same manner as described for transport distribution reconstruction measurements. The projected label positions were then used to generate diffraction limited spots as modelled by a Gaussian of width *σ*_PSF_ = 0.21·λ·*M*_ob_/*NA*_ob_ = 7.6μm as described in ^25^, hence assuming they are all in focus simultaneously. The superimposed diffraction limited spots are subsequently sampled on a 24μm pixel grid and normalized in intensity. Since the pixel-size under-samples the diffraction-limited PSF, the position of the diffraction-limited signal relative to the center of the pixel is expected to affect the accuracy of the center-detection ^22^; we therefore added a uniformly distributed random offset in both spatial coordinates to each simulated NPC in order to sample all possible sampling-biases. The simulated image of the NPC labels was subsequently either subjected to centroid extraction or used for 1D Gaussian fitting in either direction along the pixel grid. For each condition, 500 labelled NPCs were simulated. The registration error was calculated as the distance between the estimated and the ground truth positional coordinate of the pores in each direction and was found to be of zero-mean (unbiased) and roughly normally distributed. The standard deviation of the registration error *σ*_reg_ was subsequently used for comparison.

#### Pore rotation detection

The images used to determine the limitations of nuclear pore rotation detection were simulated the same way as described for the registration-error simulations, with the addition of a rotational transformation of the NPC labels along a single axis before projection. Rotation detection was either implemented by calculating the fraction of widths of fitted Gaussian functions in each spatial direction or by calculating the fraction of the length components of the principle axis found after fitting a 2D Gaussian function to the NPC signal. Cut-off values for angle-detection were based on a 95% confidence interval.

#### Simulation parameters for reconstructed radial transport distributions

Unless otherwise stated, the cargo distributions were constrained to a cylindrical central channel with radius *R* = 25nm and *l_pore_* =120nm in length. Equidistant binning of both the Cartesian projection axis Y and the radial axis *r* was used for *M* +1 bins of width *R*/*M*. Although different bin widths were simulated, all reconstructions make use of *M*=16 bins. For the sake of simplicity, sectioning along the length of the pore (*x*-axis) as shown in **Figure S1c-f** was not simulated, hence the simulations reflect a single section of length *l*. The parameters used in each simulation as well as those that were varied are summarized in the Table 1, as are the radial and angular cargo distributions used to generate the localization data.

#### Density-based comparison simulations (Figure 1)

From a biological perspective, the spatial fluctuation of particle densities within a given volume is more informative than their distribution. As a matter of fact, the numerical error of any given reconstructed transport distribution may not be a priority if the mere purpose of such a reconstruction is to distinguish between limiting cases - such as peripheral or central transport, for instance. To investigate whether 3D-SPEED would be capable of reliably making such distinctions, we simulated localization datasets as described in the previous sections according to one of four limit-case distributions: central, peripheral, bimodal and uniform (see **Table 2)**. In generating these datasets, effects of localization imprecision were introduced as described above; other compromising effects (e.g. rotation, registration, drift) were not included. The simulated localization datasets (100 replicates per condition) were subsequently transformed using the IPT, after which the resulting reconstructed radial transport density profiles were subjected to a fitting routine to determine the most likely underlying transport profile.

**Table 2.** Distributions usedfor localization dataset generation in density-based simulations.

| *Distribution name* | *Description* | *Fraction of localizations* |
| --- | --- | --- |
| Central | Gaussian: $\mu_r = 0$; $\sigma = 1.5$nm | 100% |
| Peripheral | Gaussian: $\mu_r = 23$nm; $\sigma = 1.5$nm | 100% |
| Bimodal | Gaussian: $\mu_{r,1} = 0$; $\sigma_1 = 1.5$nm | 10% |
| | Gaussian: $\mu_{r,2} = 23$nm; $\sigma_2 = 1.5$nm | 90% |
| Uniform | Uniform density $P(r) \sim r$ over $0 \leq r \leq R = 50$nm | 100% |

#### Simulation parameters for reconstructed radial transport densities

The simulation parameters were chosen to best reflect the most critical region of interest – the central channel segment of the NPC – using the dimensions as described in the original work^1^. All simulated cargo distributions were constrained to a cylindrical volume with radius *R* = 50 nm and *l_centralChannel_* = 50 nm in length. Equidistant binning of both the Cartesian projection axis Y and the radial axis *r* was used for *M*+1 bins of width *R**/**M* with *M*=10 bins. Each simulated condition consisted of a combination of one of four cylindrically symmetrical radial transport distributions (see **Table 2)**, a dataset size (100, 1000 or 10000 localizations) and a simulated localization precision *σ* (2, 4, 6 or 10nm – simulated as described before); 100 independently simulated datasets were used for reconstruction and fitting per condition. Effects of data registration, pore rotation, drift and other perturbing factors were not modelled; the simulated datasets therefore constitute a best-case scenario for the given experimental conditions.

#### Fitting routine used to identify transport density type

The reconstructed transport density profile is checked for any artefactual negative densities; if present, these are excluded from the fitting procedure (including these values or substituting them with zero before fitting resulted in lower success rates). Subsequently, a Gaussian smoothing is applied to the reconstructed density profile, the width of which was optimized for each dataset size (small datasets required more smoothing to obtain optimal results). The reconstructed density profile is then mirrored and joined to yield a symmetrical reconstructed density profile that allows central distributions to be readily fitted. A single Gaussian fit is performed, using the center radius of the bin with the highest reconstructed density value as the initial guess for the mean value *μ_1_.* If the fit residuals contain a peak value that exceeds the mean density value of the original distribution, a double Gaussian fit is performed, using the center radius of the bin with the highest residual value as the initial guess for the other mean value, *μ*_2_. The range for the fitting parameters was restricted to –*R* <*μ*_1_< *R*, –*R* <*μ*_2_< *R*, 1 ≤ *σ*_1_ ≤ *R*/2, 1 ≤*σ*_2_ ≤ *R*/2. Symmetric Gaussians were used for non-central fits (i.e. in cases where *μ*_fit_ > *σ*_fit_). If successful, the double Gaussian fit is compared to the single Gaussian fit using the following criteria:

1. The amplitude-ratio cannot be bigger than 1:5;
2. The sum of the squares of the residuals (SSR) of the double Gaussian fit needs to be smaller than that of the single Gaussian fit;
3. The double-Gaussian fit needs to be more significant than the single-Gaussian fit as measured by the following measures:

a. Adjusted R-squared value
b. Bayesian Information Criterion

If all three criteria are met, the SSR of the double Gaussian fit is compared to the SSR of a uniform distribution (constant radial density). If any of the above criteria is not met, the SSR of the single Gaussian fit is compared to the SSR of a uniform distribution. If the uniform distribution is a better fit (i.e. has the lowest SSR), the reconstructed cargo density is designated as coming from a *uniform* distribution; otherwise, the Gaussian fit is evaluated as follows:

- If the fit has a central peak, but no peripheral peak, it is designated as coming from a *central* distribution;
- If the fit has a peripheral peak, but no central peak, it is designated as coming from a *peripheral* distribution;
- If the fit has a central peak *and* a peripheral peak, it is designated as coming from a *bimodal* distribution.

For the purpose of distribution designation, a central peak is defined by *μ*_fit_ < *σ*_fit_, whereas a peripheral peak needs to satisfy *μ*_fit_ > *μ*_r,2_/2 (see **Table 2**). Only the peak(s) of the most significant Gaussian fit (single or double Gaussian, as defined by the three criteria listed earlier) are considered. All parameters for fitting (Gaussian width range, initial values), pre-fit processing (amount of smoothing, in- or exclusion of negative density values) and designation (amplitude-ratio, central/peripheral cut-off) have been optimized to maximize the success rate for identification of the ground truth distribution. Given that these distributions are not known a-priori for real datasets, the calibration of this fitting procedure represents a best-case scenario for the identifications of these types of distributions in actual datasets.

